## Supplementary material for "Environmental stiffness regulates neuronal maturation via Piezo1-mediated TTR activity": Video legends

### **Supplementary Information Videos | Titles and Legends**

**Supplementary Information Video 1 | Calcium imaging of CTRL neurons on soft gel DIV7**

Calcium imaging of wildtype control neurons on soft hydrogel. The majority of cells show peaks. Intensity is colour coded. Calcium peaks are represented by a change of colour in the corresponding cell. Video is at 4x original speed. Scale bars are 10 μm.

**Supplementary Information Video 2 | Calcium imaging of CTRL neurons on stiff gel DIV7**

Calcium imaging of wildtype control neurons on stiff hydrogel. Unlike the other conditions the cells show no peaks. Intensity is colour coded. Calcium peaks are represented by a change of colour in the corresponding cell. Video is at 4x original speed. Scale bars are 10 μm

**Supplementary Information Video 3 | Calcium imaging of P1 KD neurons on soft gel DIV7**

Calcium imaging of Piezo1 knockdown neurons on soft hydrogel. The majority of cells show peaks. Intensity is colour coded. Calcium peaks are represented by a change of colour in the corresponding cell. Video is at 4x original speed. Scale bars are 10 μm

**Supplementary Information Video 4 | Calcium imaging of P1 KD neurons on stiff gel DIV7**

Calcium imaging of Piezo1 knockdown neurons on stiff hydrogel. The majority of cells show peaks. Intensity is colour coded. Calcium peaks are represented by a change of colour in the corresponding cell. Video is at 4x original speed. Scale bars are 10 μm
