## Supplementary material for "Environmental stiffness regulates neuronal maturation via Piezo1-mediated TTR activity": Media

### Supplementary Information Methods 1 | Buffers, media, etc

#### Neuron culture media for Patch clamp and synapse staining experiments

150 ml Neurobasal
1.5 ml Glutamax
1.5 ml Penicillin-Streptomycin
3mL B-27 supplement
1.5 mL of N2

Supplemented with 2μM AraC for the first media change and 1μM AraC afterwards
filtered through top bottle filter with 220 nm pore size

#### Papain solution

2 ml HBSS
20 μl Penicillin-Streptomycin
40 units Papain
20 μl DNase I Type IV
10 μl L-Cysteine
**f**iltered through filter with 220 nm pore size

#### Ovomuccoid

50 ml of HBSS
0.5 ml Penicillin-Streptomycin
25 mg Bovine Serum Albumin
0.5 ml DNase I Type IV
50 mg Trypsin inhibitor
dissolved for 1h at 37°C

#### NB complete

150 ml Neurobasal
1.5 ml GlutaMAX
3 ml B-27
1.5 ml Penicillin-Streptomycin-Amphotericin B Mixture (PSF)
filtered with top bottle filter with 100 nm pore size

#### HBSS+

150 ml HBSS w/o Calcium, w/o Magnesium
1.5 ml PSF
filtered with top bottle filter with 100 nm pore size

#### Hibernate E+

Mix 100 ml Hibernate E
1 ml PSF
filtered with top bottle filter with 100 nm pore size

#### Magic RIPA Buffer

150 mM NaCl
1% Triton
0.5% Sodium deoxycholate
0.1% Sodium dodecyl sulfate (SDS)
50 mM Tris(Trizma base)
pH 8

#### TBS

200 mM Tris
1.37 M NaCl
pH to 7.6

#### TBST

200 mM Tris
1.37 M NaCl
0.5% Tween-20
pH to 7.6

#### Running buffer

NuPAGE™ MOPS SDS Running Buffer (20X) diluted in double distilled water = ddH_2_0

#### Complete Transfer Buffer (1L)

50 ml NuPAGE Transfer Buffer (20x)
100 ml methanol
850 ml ddH_2_O
1 ml NuPAGE antioxidant

#### 10x Marc's Modified Ringer Solution (10x MMR)

584.4 g NaCl
14.9 g KCl
20.33 g MgCl_2_6H_2_O
29.4 g CaCl_2_2H_2_O
20 ml EDTA disodium salt 0.5M
119.4 g Hepes
fill up to 10L with ddH_2_O
approximately 20 ml NaOH (used to pH to 7.8)
Buffer was aliquoted and autoclaved

#### AFM *Xenopus* Media

6.5 ml 10 x MMR
500 μl PSF
0.02 g MS222
fill up to 50 ml with ddH_2_O
pH to 7.5

#### Exposed Brain Media (EBM)

5 ml 10x MMR
20 mg MS222
500 μl PSF
fill up to 50 ml
pH to 7.4

#### Anaesthetic *Xenopus* Media

80 mg MS222
198 ml 1x MMR
2 ml PSF
pH to 7.6

#### *Xenopus* Recovery Media

1:1 Anaesthetic *Xenopus* Media, 0.1x MMR

Xenopus Rearing Media

0.1x MMR

#### PBT

PBS with 0.2% BSA and 0.1% Triton X
